# Carryover effects modulate spring phenological responses to temperature in a herbivorous insect

**DOI:** 10.64898/2026.04.01.715835

**Authors:** Sienna D. Rattigan, Léa C. Beaupère, Ben C. Sheldon, Rona Learmonth

## Abstract

1. Phenological shifts are a major ecological consequence of climate change, yet studies often focus on single life stages meaning that the potential for carryover effects between life stages remains poorly understood. Failing to account for these effects may lead to inaccurate estimates of phenological shifts, with consequences for predicted synchrony among interacting species. This is especially relevant for temperate systems where climate warming is occurring unevenly across the year.
2. Here, we investigated how temperature experienced the previous autumn and winter (during the pupal and egg stage) influences spring phenology in the winter moth (Operophtera brumata), a herbivorous insect with distinct life stages. Using 50 years of local climate data to create five experimental temperature regimes, we first quantified phenotypic plasticity in the duration and temporal variability of pupal and egg development. We then examined how timing of adult moth emergence affects timing of offspring hatching.
3. We found divergent effects of temperature on different life stages; pupal development time was shortest at intermediate temperatures while egg development time decreased linearly with increasing temperature. Furthermore, phenological shifts due to the conditions experienced by the mother were carried over to influence the phenology of her offspring. While this carryover effect was partially compensated during subsequent stages, compensation decreased under warming conditions.
4. These results refine our understanding of the sensitivity of the annual cycle of winter moth phenology to variation in temperature with potential implications for population dynamics and interspecific interactions. Overall, our findings highlight the need to consider the impacts of warming across multiple life stages so that carryover effects can be properly accounted for. Doing so will improve predictions of phenological shifts under future climates.

## Introduction

Changes in phenology (the seasonal timing of life history events) are among the most significant ecological consequences of climate change (IPCC, 2023; Parmesan and Yohe, 2003; Root et al., 2003; Thackeray et al., 2010). Cross-species analysis has shown these changes are most pronounced in primary consumers, which often show greater phenological advances with seasonal warming than other trophic levels (Thackeray et al., 2016). These systematic differences could result in widespread asynchrony between the seasonal peaks of consumer demand and resource availability. Such divergence in event timing has the potential to compromise individual fitness and potentially disrupt population dynamics in the consumer species (Lane et al., 2012; Plard et al., 2014; Simmonds et al., 2020; Visser et al., 2006), a phenomenon known as phenological mismatch (Samplonius et al., 2021).

A significant research priority is predicting if, and to what extent, phenological mismatch will occur under future climate scenarios (Visser and Gienapp, 2019). Addressing this will require a detailed understanding of how species phenology responds to key environmental variables so that we can predict future phenological change and resulting asynchrony. Phenological literature has largely focused on how changes in spring phenological events are driven by spring temperature, which has been shown to explain a large proportion of spatial and temporal variation (Badeck et al., 2004; Cohen et al., 2018; Sparks and Carey, 1995). While this has greatly advanced our understanding of phenological shifts, examining spring events in isolation, and considering only the impacts of temperature during this focal stage, may hamper efforts to predict biological responses to climate change (Williams et al., 2015). This is because phenological events are frequently part of a continuous life cycle, and conditions experienced in early life stages have the potential to influence traits and performance of later stages (Harrison et al., 2011; O’Connor et al., 2014). These delayed impacts are referred to as carryover effects.

Quantifying carryover effects by looking at multiple stages of an organism’s lifecycle is important because they can modulate phenotypic plasticity and alter phenology in ways that persist across the seasons. For example, there is growing evidence that winter conditions may shape spring phenological responses to climate change in both plants (Cook et al., 2012; Laube et al., 2014; Yuan et al., 2024) and insects (Bosch and Kemp, 2003; Chuche and Thiéry, 2009; Stålhandske et al., 2015). Failing to account for these effects may lead to over- or underestimation of phenological shifts, with consequences for predicted synchrony among interacting species. Therefore, to better understand ecological responses to climate change, we should examine phenological shifts across multiple life stages, so that potential carryover effects may be incorporated into predictive models.

Winter moths (*Operophtera brumata*) are an ideal study species for quantifying carryover effects for several reasons. Their distinct life stages show strongly plastic phenological responses to temperature (Peterson and Nilssen, 1998; Topp and Kirsten, 1991; van Asch et al., 2007). These life stages experience different seasonal conditions due to differences in annual timing and mobility (Varley et al., 1973). Moreover, as climate warming is occurring unevenly across the year (Climate Central, 2025; Renner and Zohner, 2018), temperature responses may diverge across life stages, creating potential for carryover effects to modulate plasticity. Such effects have already been observed in this system, with larval photoperiod influencing subsequent pupal development rates (Salis et al., 2017). Identifying potential-carryover effects in winter moth phenology is important because they are under strong selection to synchronise spring hatching with host tree budburst (Feeny, 1968; van Dis et al., 2023). The resulting caterpillar peak is also a key food resource for many passerine birds whose reproductive success depends on synchrony between peak chick demand and peak caterpillar abundance (Perrins, 1991; Reed et al., 2013; Simmonds et al., 2017). Carryover effects in winter moth phenology could therefore have cascading consequences across trophic levels.

In this study, we test whether carryover effects modulate phenological responses to temperature by experimentally manipulating the rearing environment of the winter moth from pupation to larval hatching (Figure 1). We quantify temperature-mediated phenotypic plasticity in the duration and temporal variability of two key life history stages (i) pupal development, defined as the number of days from pupation to adult moth emergence, and (ii) egg development, defined as the number of days from the first egg-laying event to larval hatching. Second, we examine whether the effect of temperature on the timing of adult moth emergence in winter persists to influence the timing of offspring hatching in spring. These objectives allow us to look at plasticity of individual life stages as well as carryover effects across stages.

**Figure 1:**
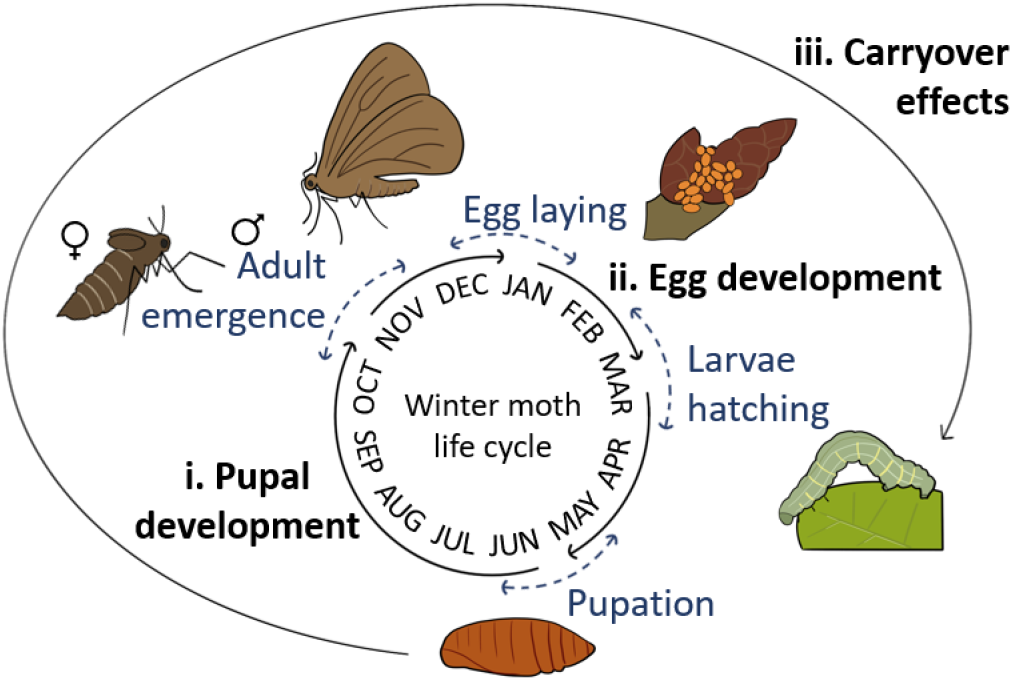
Winter moth life cycle. Adult winter moths emerge in November/December and shortly after mating brachypterous females crawl up trees to lay eggs. Eggs hatch in early spring, and larvae feed on young foliage until late May when they descend to the ground on silken threads to burrow into the soil and pupate. Pupation lasts until winter, when adults emerge and the life cycle begins again. Solid black arrows indicate the duration of each development stage and blue dashed arrows indicate the temporal variability of each stage.

## Methods

### Origin of experimental animals

We obtained our experimental animals from the surviving pool of a translocation experiment (see Learmonth et al., 2025 for full details). In November-December 2023 adult winter moths were caught with lobster pot traps on 23 oak trees covering 2.4 ha of Wytham Woods, Oxfordshire (51.77°N, 1.32°W). Captured females were kept in an outdoor storage container in individual Falcon tubes with a roll of tissue paper for oviposition. When males were captured on the same date and trap, one male was placed with each female, to increase the chance of obtaining fertilised eggs. These eggs were subsequently used in a translocation experiment, where clutches were divided into subclutches of 12 and placed into separate mesh bags. Bags were then attached around branches on rearing trees with earlier (up to 13.1 days), later (up to 17.7 days), or matching budburst phenology relative to the female’s source tree (as reported in Learmonth et al., 2025 this treatment had no detectable effects). In June 2024, the translocation experiment ended, and any winter moths surviving to the pupal stage were removed and kept in individual pots in an outdoor storage container until the start of the experiment in this study below. We tested whether the origin of these experimental animals influenced our findings, and found no evidence of any effects; this is discussed in Supplementary Information 1.

### Laboratory experiment

In July 2024, we randomly allocated pupae (n = 210) across six temperature treatments, giving 35 replicates per treatment. We created five experimental treatments in climate-controlled chambers (PHCBi MIR-154-PE) and an ambient treatment in an opaque storage box exposed to the ambient outdoor temperature at John Krebs Field Station, Oxfordshire (51.78°N, 1.31°W). The ambient treatment allowed us to account for any confounding factors, such as from humidity or air circulation, that may arise because of incubator conditions, as well as to test the response of pupae exposed to normal daily temperature fluctuations. Given that pupae develop in the soil at about 10 cm depth, where photoperiod cues are absent, we kept all six treatments in the dark for the duration of the experiment. Each pupa was kept in an individual 40 ml pot containing one of six substrate treatments, which we tested to find which maximised pupal survival by best balancing the risk of mould from excess moisture and desiccation from too little (Table S2; Figure S2).

We chose daily experimental temperature treatments to mimic the range of temperatures winter moth pupae naturally experience over their six-month development (June-December). These were calculated using Met Office daily mean temperature data for Wytham Woods at 5 km resolution (447500E, 207500N) from 1966 to 2016 (Legg et al., 2025; Parker et al., 1992). This 50-year period provided representative mean temperatures while capturing recent climate trends caused by anthropogenic emissions. First, the daily average temperature was calculated to form the ‘mean’ treatment. We then used daily standard deviation (SD) across years to create the four additional temperature gradients, each differing from the daily mean temperature by ±1.5 and 2.5 SDs to create five treatments (cold, cool, mean, warm and hot; Figure 2a).

**Figure 2:**
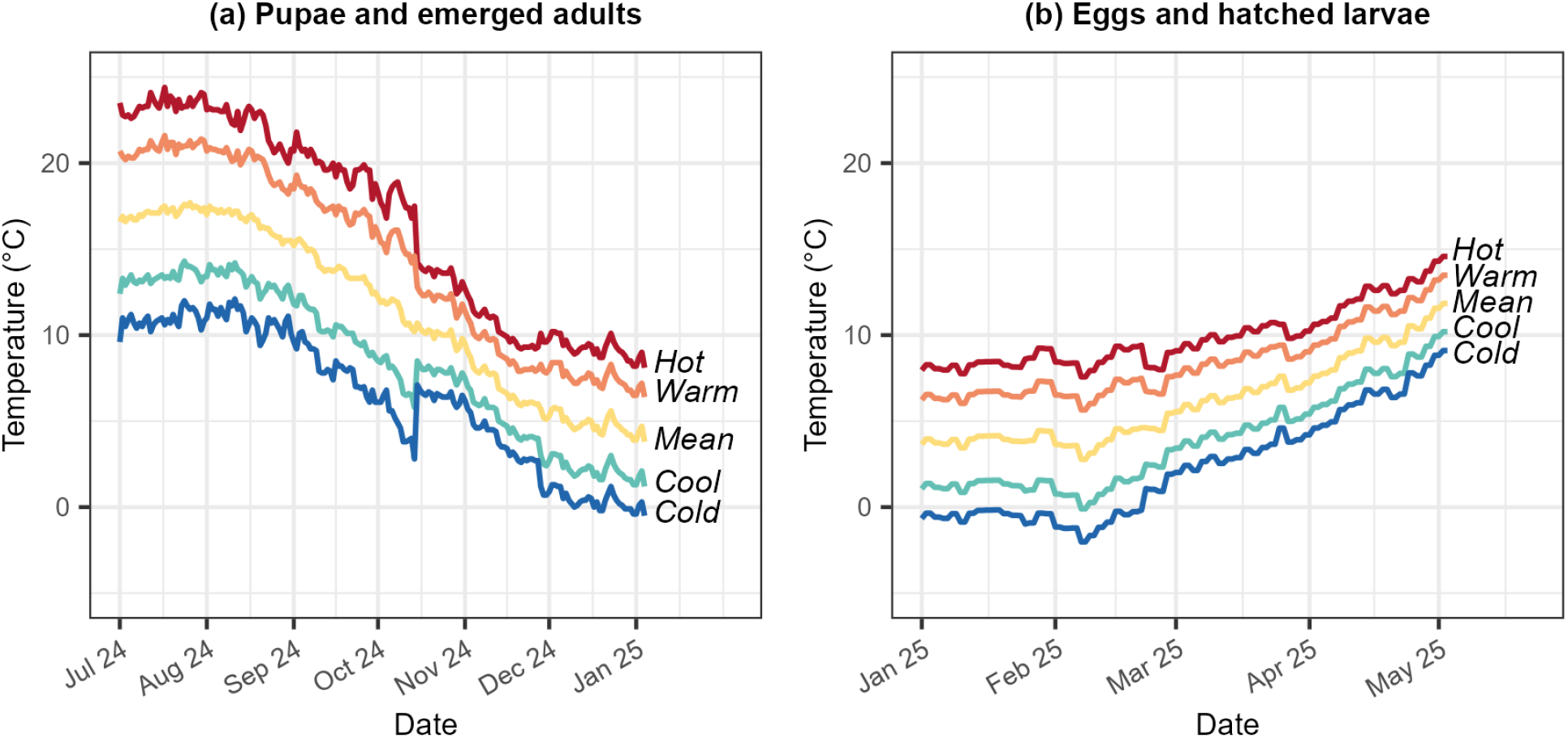
Experimental temperature profiles for **(a)** pupae and emerging adult moths and **(b)** eggs and hatching larvae. The mean treatment represents the 50-year daily mean temperature while the other four treatments represent this mean ±1.5 (cool/warm) and 2.5 (cold/hot) SDs. Daily SD was used up until October 2024 after which monthly SD is used.

Partway through the experiment, it became apparent that using daily SD would put the cold temperature treatment well below zero degrees for a prolonged period, likely reducing the chances of any moths emerging, mating or laying eggs in this treatment. To address this, the treatments were recalculated using monthly SD across years and implemented from mid-October. This was a more realistic compromise between smoothing out day-to-day fluctuations within each month while reflecting the broader year-to-year seasonal trends. In addition to capturing a range of environmental conditions, limiting temperature gradients to 1.5 and 2.5 monthly SDs above the mean reflects projected changes in mean air temperature in Wytham Woods over the next 70 years under two emissions scenarios. The warm treatment (+1.5 SDs) aligns with RCP 2.6, which assumes strong climate mitigation, limiting warming to 0.9-2.3°C by 2100 (Kendon et al., 2023). In contrast, the hot treatment (+2.5 SDs) reflects RCP 6.0, a less optimistic pathway with continued fossil fuel use, leading to 2.0-3.7°C of warming by 2100.

Since pupae were not collected as larvae, precise pupation dates for individuals were not available and we instead used the start of the experiment as a proxy, assuming that any differences were randomised across treatments. We checked all pupae for emergence every eight days between early July and September, then every four days until first emergence, then daily until early January, at which point the emergence experiment ended and remaining pupae were censored. We examined non-emerged pupae daily for mould which was removed using ethanol-dampened tissue before transferring pupae to clean pots, with the appropriate substrate treatment, to minimise re-contamination. Severely mouldy or desiccated pupae were recorded as dead and discarded.

We checked the species of emerged moths using ObsIdentify and 54/55 emergences were confirmed as winter moth, with one oak marble (*Eudemis profundana*). We then determined sex using wing morphology, as females of this species are brachypterous. This allowed us to form mated pairs, where possible, from males and females emerging from the same temperature treatment simultaneously (all matings occurred within eight days of emergence; mean = 1.96, SD = 2.47). In 3/33 cases a male from a different rearing temperature was the only available mate. Mating took place over three days in 60 ml pots with a roll of tissue paper for oviposition within the incubators. We monitored females daily to determine date of first laying event, and recorded clutch size after female death.

We then carried out a split-clutch common garden experiment across the same five experimental temperature treatments to assess maternal carryover effects. We divided each fertilised clutch (fertilized eggs can be identified by their orange coloration) with at least six eggs into six equally sized subclutches of no more than 20 eggs and placed each subclutch into separate 60 ml pots. We then randomly assigned one subclutch per female to each of the six temperature treatments, yielding 22 replicates per treatment. Given that photoperiod does not substantially affect winter moth larval hatching phenology relative to temperature (van Dis et al., 2024), we kept all eggs in climate-controlled chambers under equal light conditions which followed a monthly average light-dark regime. Eggs in the ambient treatment were exposed to natural photoperiod.

We calculated five experimental temperature treatments for eggs using the same historical temperature dataset as for the pupae. However, eggs are laid on branches and are therefore likely more exposed to temperature fluctuations throughout the day compared to pupae which are in the soil. Therefore, we interpolated daily means into daily temperature curves at two-hour resolution using the R package *chillR* (Luedeling et al., 2024). The daylength() function was used to calculate sunrise, sunset, and day length from latitude, and stack_hourly_temps() to inter-polate daily maximum, minimum, and mean temperatures to generate hourly temperatures. The average of each two-hour period was taken and repeated over two days to minimise incubator reprogramming. Nonetheless, these treatments still represent the daily mean temperatures ±1.5 and 2.5 times the monthly SD across years (Figure 2b).

Each subclutch was checked twice weekly from the middle of January. Winter moth eggs change from orange to blue-black a few days before hatching. We checked eggs daily from the first egg darkening until all eggs had hatched. At each check, a cumulative count of larvae was recorded. Any eggs still unhatched more than two weeks after the last egg in a subclutch had hatched were considered to have failed to hatch. For subclutches with more than two larvae, half-hatch date was estimated by fitting a logistic growth curve and extracting the day on which 50% of larvae had hatched. For subclutches with one or two larvae, hatching date was manually determined as the day on which the first larvae had hatched. Both estimates are referred to as ‘half-hatch date’ in analyses.

### Statistical analysis

Pupal development time was calculated as the number of days between the start of the experiment (used as a proxy for pupation date) and adult moth emergence date. The effect of temperature on pupal development time was analysed using a linear mixed-effects model (LMM) which included temperature as a continuous predictor with linear and quadratic fixed-effect terms. This required converting the categorical treatments to a continuous variable by averaging the daily temperatures from when the pupae entered the climate-controlled chambers up to the mean pupal development time in each treatment. Clutch ID was included as a random effect to account for non-independence among pupae from the same clutch.

Plasticity in pre-oviposition time, the number of days between adult moth emergence and the first egg-laying event, was explored but a small sample size limited the inference possible (Figure S3). Furthermore, while 29 females laid eggs, only 22 clutches were suitable to use in subsequent experiments and analyses. These 22 clutches were fertilised eggs from females that had been mated once to a male reared in the same temperature treatment.

Egg development time was calculated as the number of days between the first egg laying event of the clutch and the half-hatch date of the subclutch. The effect of temperature on egg development time was analysed using a LMM which included temperature as a continuous linear fixed-effect term. This also required converting the categorical temperature treatments to a continuous variable by averaging the daily temperatures from the start of the split-clutch experiment to the mean egg development time in each treatment. This model included maternal ID as a random effect to account for non-independence among larvae hatching from eggs laid by the same female.

In addition to plasticity in life cycle timing, we also analysed adult moth survival and reproduction to assess how plasticity in phenology may influence population dynamics. We applied a Kaplan-Meier survival analysis to model pupal emergence success and used a log-rank test to compare survival across temperature treatments (for the purposes of this model we classified moths in pupa as ‘surviving’ and those that had emerged as ‘non-surviving’). We then analysed clutch size using a generalised linear model which included temperature as a continuous predictor with linear and quadratic fixed-effect terms. Females which did not lay any eggs were included in this model as clutch sizes of zero.

Lastly, to analyse phenological carry-over effects we used LMMs to examine how maternal emergence date influences (i) subclutch half-hatch date and (ii) egg development time. Both models included egg temperature treatment as a categorical fixed-effect covariate to account for variation in the temperature experienced by subclutches. For full details of all models see Table S4.

We conducted all statistical analyses using R version 4.5.2 (R Core Team, 2026). We included quadratic or interaction terms only where they significantly improved model fit in a likelihood ratio test. As an overall test of key predictors, full models were compared to null models that were identical except lacking the main effects. Various combinations of random effects were fitted in all models including source tree of adult moths in 2023, rearing tree in the 2024 translocation experiment, original clutch identity, and maternal identity in the next generation. Random effects were only retained when the model was not overfit and when a significant proportion of the variance was explained. We checked all model diagnostics and confirmed that they met the assumptions required.

## Results

Of 210 pupae, 54 emerged, with development times between 127 and 187 days (mean = 153, SD = 14). Amongst these were 32 females, 29 of which laid eggs with clutch sizes between one and 272 eggs (mean = 117, SD = 92). We used 22 of these clutches for further experimentation and analyses as they were fertilised and came from females that had been mated once to a male reared in the same temperature treatment. Hatching was observed in 86% of the 132 subclutches with egg development times between 102 and 186 days (mean = 136, SD = 19).

### Effect of temperature on pupal development time

Temperature had a significant nonlinear effect on pupal development time, with adult moth emergence delayed at extreme temperatures relative to intermediate ones (Figure 3). Pupal development time initially decreased with increasing temperature (effect of temperature: -14.01 ± 3.91, p < 0.001), however this trend reversed at warmer temperatures (effect of temperature2: +0.99 ± 0.16, p < 0.001). Expected pupal development time was shortest (∼143 days) at temperatures averaging 12.2°C pre-emergence. Although the ambient treatment experienced greater temperature variation, pupal development time matched expectations from the experimental treatments, as ambient temperatures over this period in 2024 were comparable to those in the mean treatment (Table S5; Figure S5).

**Figure 3:**
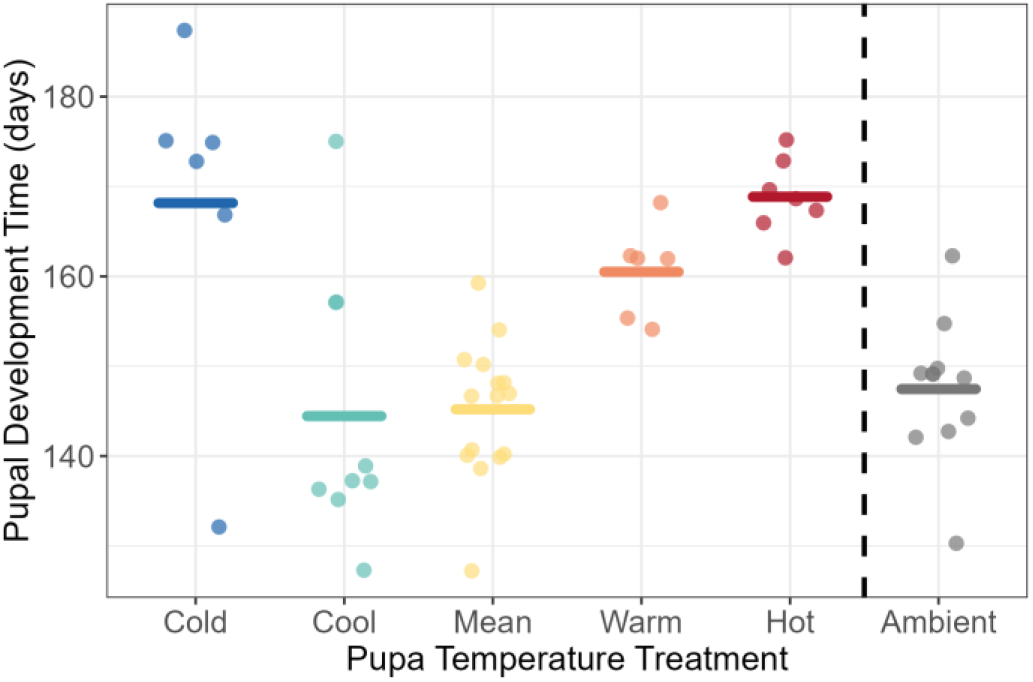
The effect of temperature treatment on pupal development time. Each coloured point represents the number of days between the start of the experiment (used as a proxy for pupation date) and the emergence date of an individual adult moth. Horizontal bars indicate treatment means and the dashed vertical line separates the experimental treatments from the ambient.

### Effect of temperature on adult moth fitness

The Kaplan-Meier plot (Figure 4a) shows the proportion of pupae yet to emerge over time for the six temperature treatments. A total of 84 pupae (40%) were censored, that is, they had not been classed as inviable at the end of the experiment, but no moth had emerged by day 188 of pupal development. A log-rank test confirmed that temperature has a significant effect on pupal survival to the adult stage (χ^2^ = 14.6, df = 5, p = 0.01). Notably, pupal development times were longest in the hot, warm and cold treatments while emergence occurred earlier and more frequently in the mean, ambient and cool treatments. These results imply temperature significantly affects pupal emergence success, although it is possible that some additional moths might have emerged in the extreme temperatures, had the experiment continued for longer.

**Figure 4:**
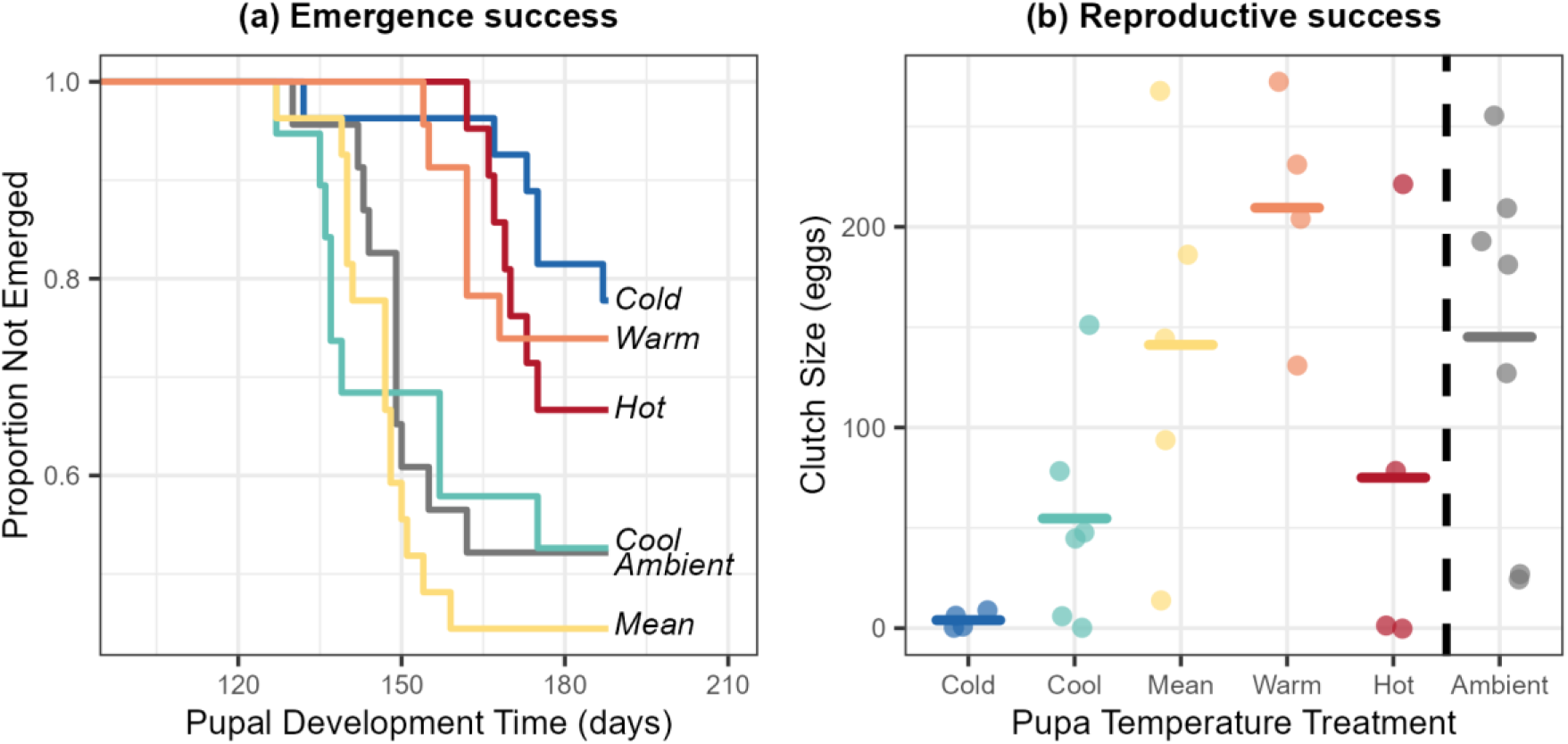
The effect of pupal temperature on adult moth fitness. **(a)** Kaplan-Meier survival curves showing pupal survival probability over time. **(b)** The effect of pupal temperature treatment on clutch size. Each point represents the number of eggs laid by an individual female adult moth (n = 29). Horizontal bars indicate treatment means and the dashed vertical line separates the experimental treatments from the ambient. Colours show the six temperature treatments in both plots.

We also found that clutch size initially increased with temperature (Figure 4b; effect of temperature: +2.12 ± 0.63, p < 0.001) but this trend reversed at higher temperatures (effect of temper-ature2: -0.07 ± 0.03, p = 0.033). Expected clutch size was maximised (∼180 eggs) by pre-emergence temperatures of 14.5°C. Again, clutch size in the ambient was comparable to the mean, despite its exposure to greater temperature variation (Table S5; Figure S5).

### Effect of temperature on egg development time

Temperature had a significant linear effect on egg development time with advanced larval hatching at hotter temperatures (Figure 5). Average egg development time in the mean temperature treatment was ∼141 days (SE = 2.045). Relative to the mean, eggs in the warm and hot treatments developed ∼16 (SE = 1.314, p < 0.001) and ∼28 (SE = 1.307, p < 0.001) days faster, respectively, whereas eggs in the cool and cold treatments developed ∼12 (SE = 1.310, p < 0.001) and ∼20 (SE = 1.314, p < 0.001) days slower. Although the ambient treatment experienced greater temperature variation, egg development time matched expectations from the experimental treatments, as ambient temperatures during this period in 2025 were comparable to the warm treatment (Table S5; Figure S5).

**Figure 5:**
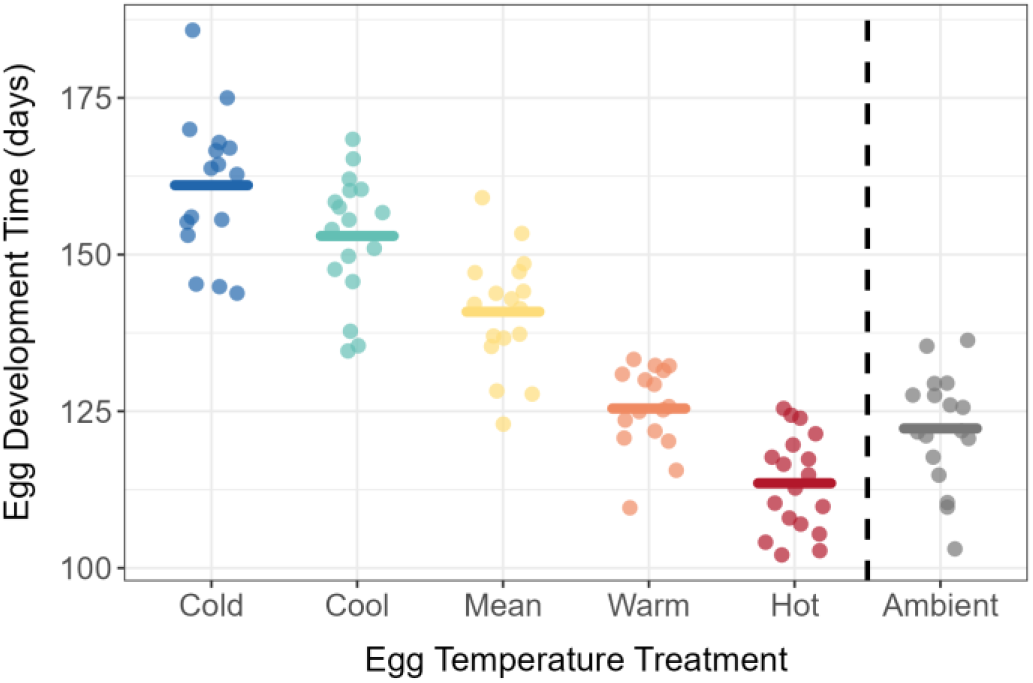
The effect of temperature treatment on egg development time. Each coloured point represents the number of days between the first egg-laying event and half-hatch date of each subclutch. Horizontal bars indicate treatment means and the dashed vertical line separates the experimental treatments from the ambient.

### Effect of maternal emergence date on egg development time

Females which emerged later in winter laid eggs that hatched later the following spring (est: +0.27 ± 0.09, p = 0.003). This relationship held across the different egg temperature treatments such that later emerging females always produced offspring that hatched later. However, the slope of this maternal effect is substantially less than one, meaning delays in emergence are not directly translated into equally large delays in larval hatching.

We further found that the eggs of later emerging females had shorter development times than those emerging earlier (Figure 6; F(1, 11.56) = 96.01, p < 0.001). Additionally, the strength of this relationship varied across egg temperature treatments (F_(4, 51.52)_ = 8.34, p < 0.001). For every one-day delay in maternal emergence, egg development time in the mean treatment was shortened by ∼0.73 days on average (SE = 0.09, p < 0.001). By comparing the slopes between the experimental and mean treatments, we found that this effect decreased under higher temperatures, with egg development time in the warm and hot treatments shortening by ∼0.45 (SE = 0.10, p = 0.005) and ∼0.46 (SE = 0.09, p = 0.007) days respectively, per day emergence was delayed. Relative to the mean treatment, egg hatching time advanced more rapidly with maternal emergence date in colder temperatures, with egg development time in the cool and cold treatments shortening by ∼0.80 (SE = 0.09, p = 0.432) and ∼0.87 (SE = 0.10, p = 0.153) days respectively, per day emergence was delayed. However, the greater delay in the colder treatments compared to the mean treatment was not statistically significant.

**Figure 6:**
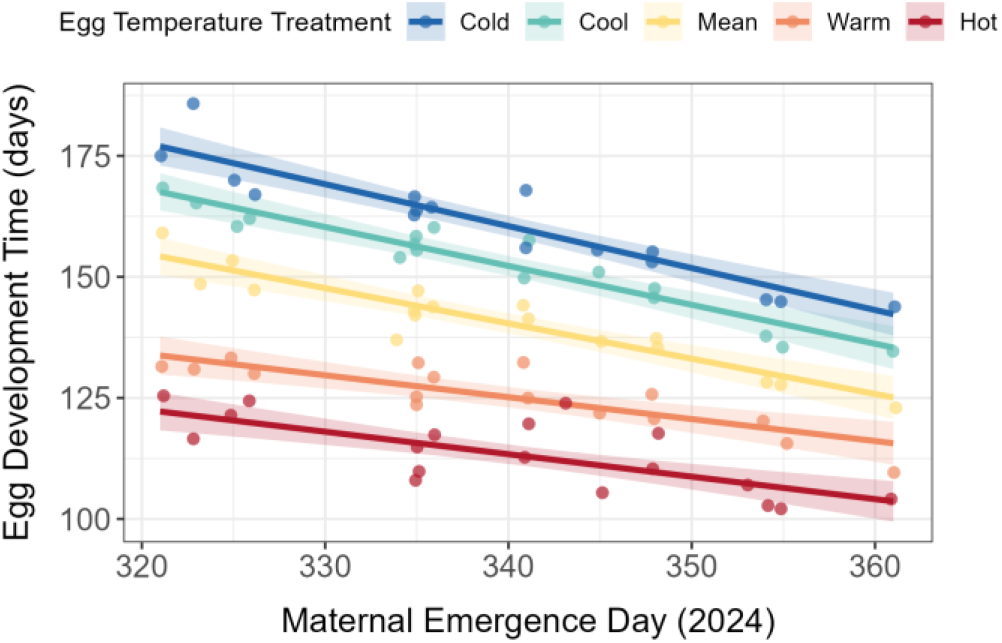
The effect of maternal emergence date on offspring’s egg development time. Points represent subclutches so those aligned in the x-axis are from the same mother. Lines show the model prediction with a 95% confidence interval represented by the shaded ribbon. Both are coloured by egg temperature treatment.

## Discussion

We experimentally tested whether carryover effects modulate phenological responses to temperature in the winter moth. We found divergent effects of temperature on different life stages; pupal development time was shortest at intermediate temperatures (Figure 3) while egg development time decreased linearly with increasing temperature (Figure 5). We further found that eggs laid by later-emerging females hatched later but had shorter development times compared to eggs laid by earlier-emerging individuals (Figure 6). Our results suggest that while temperature-mediated plasticity in adult moth emergence date is carried over to influence offspring hatching, delays are partially compensated later in the life cycle during egg development. However, the extent of this compensation varied depending on egg temperature treatment, which raises questions about the ability of winter moth populations to maintain synchrony with host trees under rapid climate change.

We found that pupal emergence was delayed at extreme temperatures relative to intermediate ones. Notably, adult winter moths reared in the cold and hot treatments as pupae emerged on average 25 days later than those in the mean. This agrees with earlier work showing that extreme low or high temperatures both delay winter moth emergence (Peterson and Nilssen, 1998; Topp and Kirsten, 1991). While previous studies have used constant temperature treatments, we used regimes that changed temperature daily and were calculated based on the 50-year local average. By doing so, we provide a more realistic representation of the natural conditions occurring during winter moth pupation, which takes place over several months. This is supported by field data from live winter moth females sampled in Wytham Woods in 2024 by Learmonth et al. (2025). The average capture date of these moths (taken as a proxy for their emergence date) was 5^th^ December, which closely matches the average emergence date of moths in the mean and ambient treatments of this study which was 28^th^ November.

A possible explanation for the non-linear pattern between pupal development and temperature is that intermediate temperatures fall within the thermal optimum for winter moth pupation, resulting in the fastest developmental rate. Temperatures below this optimum may slow down metabolism (Angilletta, 2009; Gillooly et al., 2001), while temperatures above this optimum may increase metabolism to such an extent that organisms experience oxidative stress due to the energy imbalance (Clarke, 2003; Kuo et al., 2023). Both extremes could result in a slower pupal developmental rate, delaying winter moth emergence in autumn/winter. Alternatively, pupae may have both a chilling and heating requirement to trigger emergence with intermediate temperatures providing the right balance of both cues (Forrest and Miller-Rushing, 2010). In this case, colder temperatures fail to accumulate enough heat units, while hotter temperatures bypass the chilling requirement altogether explaining the delayed emergence in both treatment extremes.

Adult moths that emerged were mated within temperature treatments and allowed to lay eggs which were used in a split-clutch experiment to isolate environmental and genetic effects. We found that eggs reared in the hot temperature treatment hatched on average 48 days earlier than those in the cold, giving a climate sensitivity of ∼7 days advance per °C increase in temperature. This finding confirms previous experimental (Buse and Good, 1996; Learmonth et al., 2025; van Dis et al., 2024) and long-term observational studies (Salis et al., 2016; Visser and Holleman, 2001). While there was some variation between subclutches within each temperature treatment, we found this was largely explained by the identity of the mother which is consistent with the known genetic component explaining variation in this trait (van Asch et al., 2007).

Combining our findings on pupal and egg development to look across the life cycle revealed evidence of carryover effects. Females exposed to higher temperatures as pupae emerged later and laid eggs that hatched later. Although carryover effects of temperature have not previously been quantified in the winter moth’s life cycle, similar effects driven by photoperiod have been reported in other lifecycle stage transitions. Notably, Salis et al. (2017) found that winter moth females receiving an early-season photoperiod as larvae laid eggs that took less time to develop than those receiving a late-season photoperiod treatment. As here, the effect size of these carryover effects depended on the temperature treatment experienced by the eggs. Building on this finding, future work could test if carryover effects modulate phenological responses to temperature at other lifecycle transitions, such as from larval hatching to pupation and pupation to adult moth emergence.

The importance of finding parent-offspring carryover effects from autumn/winter emergence to spring hatching can be illustrated by considering two contrasting scenarios. First, if we consider only the effects of spring temperature on egg development, our work (and previous studies Buse and Good, 1996; van Dis et al., 2024), predict that warmer conditions will cause winter moth eggs to hatch earlier in spring. Earlier hatching would increase the risk of asynchrony with host trees budburst (van Asch et al., 2013; Visser and Holleman, 2001), potentially leading to severe fitness consequences through larval starvation (van Dis et al., 2023). However, if we also account for the transgenerational effects of temperature experienced the previous season, by incorporating maternal emergence date into analyses, predictions change. Carryover effects from the pupal stage may instead lead to delayed larval hatching under the same warming scenario. Delayed hatching could likewise generate asynchrony with host trees, reducing larval performance due to increased leaf tannin concentrations (Feeny, 1968). The reversal in the direction of the winter moth’s phenological shift after accounting for maternal emergence date highlights the importance of carryover effects in influencing the synchrony of interactions in this and potentially other similar systems.

In addition, however, we also found that eggs from later-emerging females took less time to develop in comparison to eggs from those emerging earlier. This suggests a compensatory maternal effect, where the conditions the mother experienced influenced her offspring’s development time in a way that partially offsets the carryover effect associated with delayed emergence (Bernardo, 1996). Such maternal effects can occur when mothers allocate non-genetic components such as hormones and epigenetic markers via eggs, which influence the offspring’s developmental trajectory (Mousseau and Fox, 1998). Whether or not the maternal effects found in this study are adaptive requires further exploration into their links to offspring fitness components, for example, larval growth rate and survival to pupation.

While compensatory effects that shortened egg development time may partially buffer the extent of delays that are carried over, we further found that the strength of compensation decreased when eggs experienced higher temperatures. Specifically, the delay of larval hatching from later-emerging females was significantly greater for eggs in the hot and warm treatments than the colder treatments. Therefore, climate warming may mean winter moths are less able to buffer carryover effects from previous life stages going beyond simple shifts toward earlier seasonal activity (Forrest, 2016), which could increase the likelihood and severity of mismatches. To understand why compensation may decrease under higher temperatures in this system, future work could use a greater range of warmer temperatures to test whether winter moth eggs have a minimum development threshold that has to be met which limits the strength of compensation.

To summarise, we show that divergent temperature responses at different stages and carryover effects between stages may alter insect phenology, and synchrony with host species, in unexpected ways. Our findings refine understanding of annual variation in winter moth phenology and highlight the need to consider warming effects across multiple life stages. For example, future studies should consider the impact of warming in winter and spring rather than examining spring events in isolation. Accounting for carryover effects in this way may improve predictions of asynchrony under future climates, which is vital for identifying populations potentially at risk of phenological mismatch.

## Acknowledgements

We thank Sam Croft and Lucy Morley for help with field collection and David López-Idiáquez for help in the lab. This work was supported by grants from the UKRI Frontiers award EP/X024520/1 to BCS.

## Author contributions

Conceptualisation: SDR, BCS, RL; Data collection: SDR, LCB; Methodology: SDR, RL; Formal Analysis: SDR; Visualization: SDR; Interpretation of results: SDR, BCS, RL; Writing - Original Draft: SDR; Writing -Review & Editing: SDR, LCB, BCS, RL.

## Data availability statement

Code and data to reproduce all analyses conducted in this paper can be found at https://github.com/siennarattigan/carryover_effects.git

