## Supplementary information for "Carryover effects modulate spring phenological responses to temperature in a herbivorous insect"

### 1. *Origin of experimental animals*

We tested whether the origin of our experimental animals influenced our findings. We found that there was no significant effect when translocation treatment or original clutch ID in 2023 was included as a fixed or random effect. Furthermore, any differences we did not detect were presumably randomised across treatments when surviving pupae were assigned to temperature treatments.

### 2. *Pupal substrate treatments*

We tested a range of six substrate treatments (Table S2). We found that survival to adult moth emergence was significantly higher for treatments with vermiculite on top of cotton wool and that were sprayed with distilled water every 8-16 days ($\chi^2$ = 42.04, df = 5, p < 0.001). This is supported by Figure S2 which shows that ~75% of pupae emerged in substrate treatments A and B. This result should provide a new protocol for rearing winter moth pupae which will increase sample sizes in future work.

| **Treatment** | **Conditions** |
|---|---|
| A | Vermiculite, cotton wool, spray with distilled water every eight days |
| B | Vermiculite, cotton wool, spray with distilled water every 16 days |
| C | Vermiculite, spray with distilled water every eight days |
| D | Vermiculite, spray with distilled water every 16 days |
| E | Vermiculite only |
| F | Vermiculite only |

**Table S2:** Six substrate treatments for pupae.

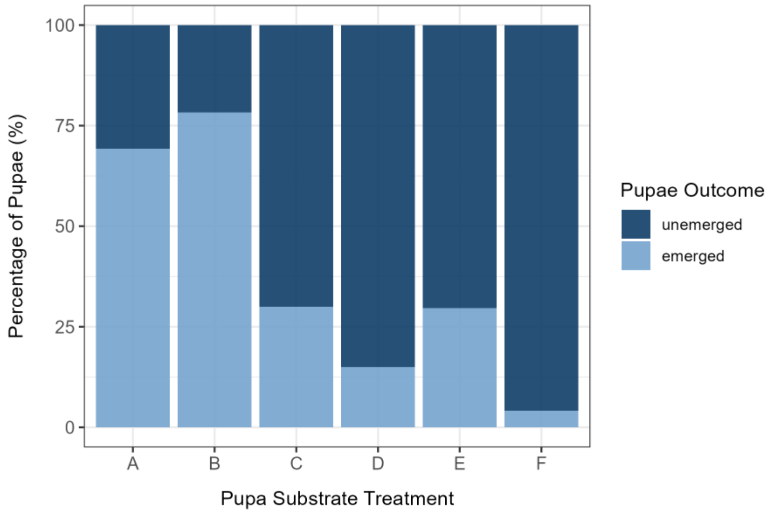


**Figure S2:** Pupal emergence success across substrate treatments. Colours show outcome by day 188 of pupal development.

3. *Pre-oviposition time*

Of 33 female moths that emerged, 22 laid eggs and were mated once to a male in the same treatment with pre-oviposition time ranging from 2 to 14 days (mean = 7, SD = 3). Temperature treatment had a significant effect on pre-oviposition time ($F_{(2,18)}$ = 3.566, $R^2$ = 0.204, p = 0.050; Figure S3). The linear component of the model was significant (effect of temperature: -2.21 ± 0.80, p = 0.013), indicating that pre-oviposition time initially decreases with increasing temperature. The quadratic component of the model was also significant (effect of temperature$^2$: +0.19 ± 0.08, p = 0.027), showing that this linear trend reverses with further warming. The minimum from the model prediction is 5.71°C, representing the average seven-day temperature pre-emergence at which pre-oviposition time is expected to be fastest (~6 days after emergence)

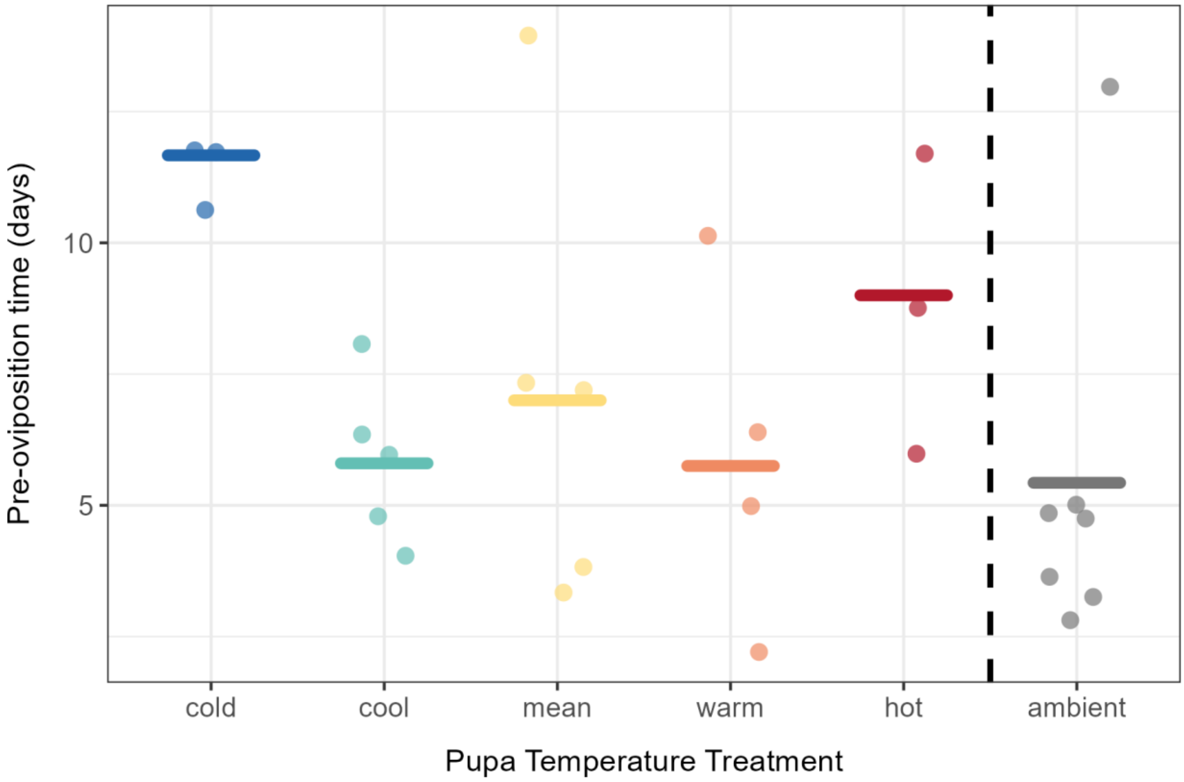


**Figure S3:** The effect of temperature treatment on pre-oviposition time. Each coloured point represents the number of days between adult moth emergence and the first egg-laying event of each female. Horizontal bars indicate treatment means and the dashed vertical line separates the experimental treatments from the ambient.

*4. Model structure*

| **Dependent variable** | **Sample Size** | **Fixed effects** | **Random effects** | **Model type** |
|---|---|---|---|---|
| Pupal development time | Emerged moths in five experimental treatments: n=43 | Continuous pupal temperature treatment and its square | Clutch ID | LMM |
| Clutch size | Clutches of greater than six fertilised eggs (excluding females mated to a male from a different treatment): n=22 | Continuous pupal temperature treatment and its square | NA | GLM (with negative binomial error distribution and log link function) |
| Egg development time | Subclutches with hatched larvae (excluding females mated twice or to a male from a different treatment): n=53 | Continuous egg temperature treatment | Maternal ID | LMM |
| Larval half-hatch date | Subclutches with hatched larvae (excluding females mated twice or to a male from a different treatment): n=53 | Adult moth emergence date, categorical egg temperature treatment and their interaction | Maternal ID | LMM |
| Egg development time | Subclutches with hatched larvae (excluding females mated twice or to a male from a different treatment): n=53 | Adult moth emergence date, categorical egg temperature treatment and their interaction | Maternal ID | LMM |

**Table S4:** Model structures.

*5. Mean vs ambient treatments*

Comparing the temperature profiles for the ambient and mean treatments reveals the expected greater variability under ambient conditions (Figure S5). Nevertheless, the average temperature experienced by the pupae did not differ between these two treatments, nor did pupal and adult moth phenology (Table S5a). However, the average temperature experienced by eggs in the ambient was significantly higher, by ~1.6°C, than the mean treatment due to a very warm March/April in 2025, putting it closer to the warm experimental treatment. This is reflected in a significantly earlier larval half-hatch date and shorter egg development time in the ambient when compared to the mean (Table S5b).

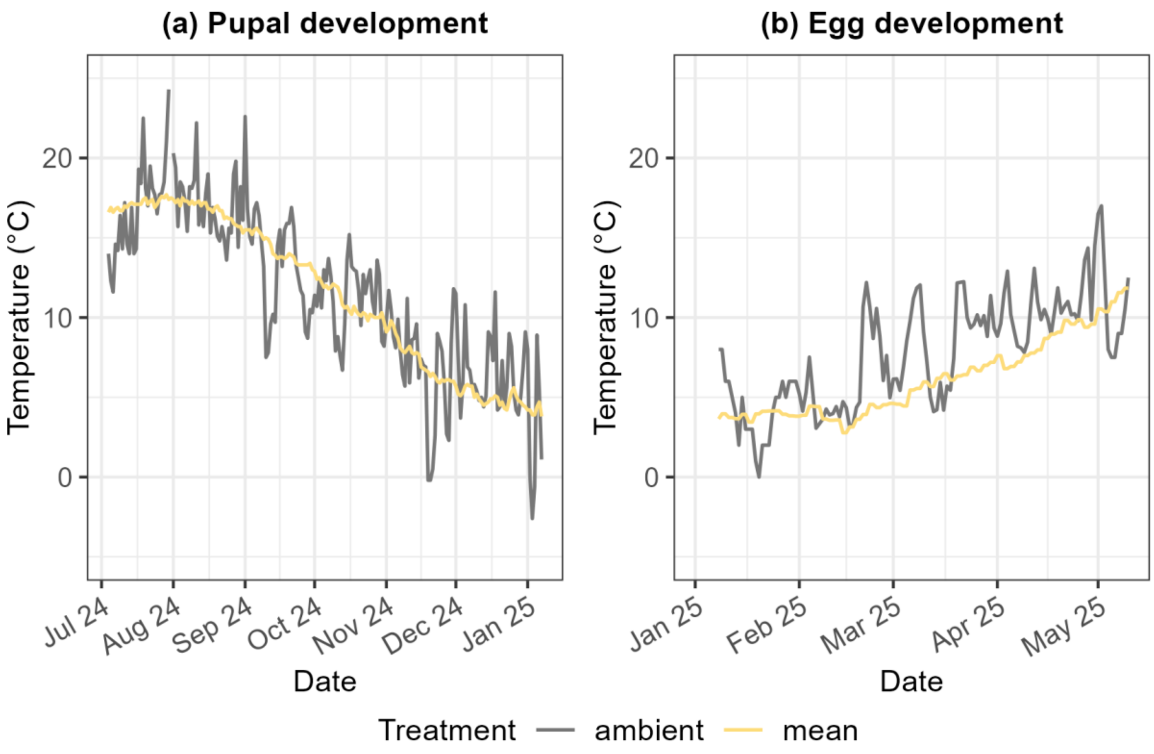


**Figure S5:** Daily temperature profiles for **(a)** pupae and **(b)** eggs in the mean and ambient treatments.

| | | Mean | Ambient | t-value | p-value |
|---|---|---|---|---|---|
| **(a) Pupae & Adults** | Average temperature (°C) | 11.54 | 11.68 | -0.25 | 0.80 |
| | Pupal development time (days) | 145.2 | 147.5 | -0.72 | 0.48 |
| | Pre-oviposition time (days) | 7.0 | 4.1 | 1.13 | 0.34 |
| | Clutch size (eggs) | 141.2 | 145.1 | -0.07 | 0.94 |
| **(b) Eggs & Larvae** | Average temperature (°C) | 5.68 | 7.33 | -4.97 | <0.001 |
| | Larval half-hatch day (Julian) | 114.2 | 98.5 | 10.88 | <0.001 |
| | Egg development time (days) | 139.6 | 120.5 | 5.72 | <0.001 |

**Table S5:** Results of t-tests comparing temperature experienced and performance of **(a)** pupae and adult moths and **(b)** eggs and larvae in the mean and ambient treatments.
